# Assessment of ferroptosis inducers and Nrf2 inhibitors as radiosensitisers in 2D and 3D breast cancer cell cultures

**DOI:** 10.1101/2023.12.08.570735

**Authors:** AA Alzufairi, C Souilhol, N Jordan-Mahy, NA Cross

## Abstract

Ferroptosis is a form of programmed cell death that is modulated in some cancer cells as a pro-survival mechanism. Induction of ferroptosis is a potential anti-cancer strategy, and enhancement of ferroptosis has the potential to current anti-tumour mechanisms. In this study, we assessed the effect of the ferroptosis inducers Erastin, RSL-3 and FIN-56 on the radiosensitivity of MCF-7 and MDA-MB-231 breast cancer cell lines in 2D and 3D cell culture. Since some tumours modulate ferroptosis via increased Nrf2 production, and MCF-7 and MDA-MB-231 both produce Nrf2 protein, we also assessed the effects of the Nrf2 inhibitor ML385 on radiosensitivity. MDA-MB-231 was highly sensitive to all ferroptosis inducers, and ferroptosis was reversed by the ferroptosis inhibitors Ferrostatin-1, Liproxstatin-1 and Deferoxamine. MCF-7 was resistant to all ferroptosis inducers. MDA-MB-231 and MCF-7 cells were sensitive to irradiation in 2D cell culture but resistant in 3D alginate spheroids. There was also no robust enhancement to irradiation effects with ferroptosis inducers or ML385 in 2D or 3D cell culture or colony formation assays. Ferroptosis inducers did, however, show heterogeneous responses in 3D cell culture. These studies suggest targeting ferroptosis does not induce enhancement of ferroptotic cell death in breast cancer cells.

## Introduction

Ferroptosis is an iron-dependent form of cell death that is characterised by the accumulation of reactive oxygen species (ROS) which induces lipid peroxidation causing morphological features differently from other forms of programmed cell death.^1^ Inducers of ferroptosis typically target either glutathione peroxidase 4 (GPX4), which protect against lipid peroxidation and is reliant on glutathione, or target glutathione biosynthesis.^2^ For example, Erastin directly inhibits system Xc, which is the cystine/glutamate antiporter crucial for subsequent glutathione biosynthesis from cysteine and glutamate.^3^ Likewise, Ras Synthetic Lethal-3 (RSL3) binds and inactivates GPX4, leading to accumulation of lipid peroxidation, which leads to ferroptotic cell death.^4^ FIN56 induces GPX4 degradation although the mechanism is unclear.^5^ Ferroptosis can be inhibited using a panel of small molecule ferroptosis inhibitors. Deferoxamine is a classic iron-chelating ferroptosis inhibitor that can be used to prevent ferroptosis and damage to normal cells and tissues.^67^ Liproxstatin-1 and Ferrostatin-1 are potent inhibitors of lipid peroxidation and are able to reverse ferroptosis in GPX4^−/−^ cells and in response to ferroptosis inducers.^8^ Liproxstatin-1 also significantly increases nuclear factor erythroid 2-related factor 2 (Nrf2), the master regulator of the antioxidant response, which helps to promote cell survival.^9^ Both Erastin and RSL3 inhibit the ferroptosis pathway in cancer cells by disturbing the redox homeostasis and allowing the iron-independent accumulation of lethal ROS.^1^ Ferroptosis can be induced by blocking system Xc-with small molecule ferroptosis inhibitors such as Erastin. Ferroptosis can also be induced by inhibition of GPX4 stability or activity using FIN56 or RSL3 respectively.^10^ Erastin reduces cellular GSH levels by inhibiting system Xc-directly, reducing cystine and glutamate uptake which are crucial for glutathione production, rendering cells susceptible to lipid peroxidation. RSL3 induces ferroptosis by targeting GPX4 directly.^11^

Irradiation is known to induce cell death by generating ROS, depleting glutathione, and activating targets such as acyl-CoA synthetase long-chain family member 4 (ACSL4), which alters cellular lipid composition.^12^ Furthermore, ferroptosis is a key mediator of radiation-induced injury, and ferroptosis inhibition mitigates radiation-induced injury.^12^ Recent studies have identified GPX4 inhibitors that potentiate irradiation responses,^13^ and ferroptosis is as a key mechanism of irradiation-induced cell death in colon cancer cells, albeit less potently in hypoxia.^14^ Furthermore, ferroptosis inhibitors can reverse radiation-induced cell death, whereas the ferroptosis inducers RSL3 enhance it.^15^ Therefore, we hypothesised that ferroptosis inducers would enhanced radiotherapy responses in breast cancer cells, and we have expanded these studies to 3D cell] culture models, which exhibit considerable hypoxia. Furthermore, Nrf2 is a master regulator of the antioxidant response and is intrinsically linked in to modulating ferroptosis signalling.^16^ It is sequestered by Kelch-like ECH-associated protein 1 (Keap-1), and in the presence of reactive oxygen species, it is released.^17^ Furthermore, Nrf2 is a prerequisite for 3D spheroid formation, and the knockdown of Nrf2, prevents spheroid formation; presumably due to the loss of an antioxidant response.^16^ The specific role of Nrf2 in mediating radio-responses in 3D cell culture is not known. Nrf2 is known to respond to irradiation and DNA damage, which is mediated by an increase in glutathione during irradiation exposure.^18^ Furthermore, radio-resistance in breast cancer stem cells have been linked to the activation of the Keap1-Nrf2 axis.^19^ Additionally, Nrf2-inibition using ML385, sensitised breast cancer stem cells to ionising radiation, showing that irradiation-induced ROS can activate Nrf2, counteracting the irradiation-induced damage ^20^ and the Nrf2-inhibitor ML385 promoted ferroptosis in eosophageal squamous cell carcinoma cell lines.^21^ Therefore, Nrf2 inhibition might be a potent radio-sensitising strategy.^20^ Although studies have focussed on cancer stem cells in breast cancer, studies have not directly assessed the cellular responses to irradiation in tumour spheroids. It is therefore hypothesised that ferroptosis inducers increase death after irradiation of both 2D and 3D breast cancer cell culture models, and that this is reversed by ferroptosis inhibitors.

## Materials and methods

### Cell culture and treatment of cells with ferroptosis inducers and inhibitors

MCF-7 and MDA-MB-231 (ECACC) were cultured in Dulbecco’s Modified Eagle Medium (DMEM) (Lonza, Manchester, UK), supplemented with 10% (v/v) heat-inactivated foetal bovine serum (FBS) (Lonza) and 1% of 100 IU penicillin and 100 µg/ml streptomycin (ThermoFisher Scientific, Altrincham, UK) incubated at 37°C under 5% CO_2_. For cell treatments in 2D cell culture, cells were plated at 1×10^4^ cells/well in a 96-well plate and left to adhere for 24 hours prior to treatment. Cells were treated with either RSL3 (Sigma, Poole, UK), Erastin (Selleck Chem, Ely, UK) or FIN56 (Sigma) at 0-10 µM in the presence or absence of the ferroptosis inhibitors Deferoxamine (Selleck Chemical company Ely, UK) at 10 µM, Liproxstatin-1 (Sigma) at 1 µM or Ferrostatin-1 (Sigma) at 1 µM. For ML385 treatments, cells were treated with 10 µM ML385 (Sigma). All treatments and controls contained a final concentration of 0.2% Dimethyl sulfoxide (DMSO) as a vehicle control. All treatments were carried out in triplicate in three technical repeats.

### Irradiation of cells

Cells were either plated in 96 well plates at a cell density of 1×10^5^ cells/ml or for colony formation assays, prepared at 1×10^5^ cells/ml. Cells were exposed to a ^137^Cs source delivering 1 Gy per 28 seconds at 0.6MeV. After irradiation, cells were either treated with ferroptosis inducers and/or ML385 or plated out for colony formation assays. In all experiments, a control plate or tube of cells was mock-irradiated.

### Assessment of cell death using Hoechst 33342 and propidium iodide staining

After cell treatments, cells were stained with Hoechst 33342 (Sigma) and PI (Sigma) at a final concentration 10 µg/ml for 30 minutes, examined using an IX70 fluorescence microscope (Olympus) and images captured using Cell-F software (Olympus). Cells were counted manually, and the percentage of cell death was calculated based on duplicate representative fields of view each containing at least 100 cells.

### CellTiter-Glo® luminescent cell viability assay

Cells were seeded in white 96-well plates (Fisher Scientific) at 1×10^4^ cell per well and treated with each ferroptosis inducers (RSL3, Erastin, FIN56) alongside a 0.2% (v/v) DMSO vehicle control for 24 hours. After treatments, 25 µl of CellTiter-Glo® reagent from CellTiter-Glo® luminescent cell viability assay kit (Promega, Southampton, UK) was added per well and mixed and incubated at room temperature (RT) for 10 minutes and measured using a CLARIOstar plate reader (BMG Labtech, Ortenberg, Germany). The average from three luminescence measurements was calculated and all treated cells were normalized to the vehicle controls, which was assigned a 100% ATP activity in three independent experiments.

### 3D Alginate cell culture assay

MDA-MB-231 and MCF-7 cultured cells were resuspended at 1×10^6^ in 0.15M NaCl and resuspended in 1mL of a sterile 1.2% (w/v in Saline) sodium alginate (Sigma) and dropped into 15-20 ml of sterile 0.2M CaCl_2_ in a 50ml falcon tube using a 21G needle (Sigma-Aldrich). This solution was then incubated at 37°C for 5 minutes. Beads were then washed twice in 15ml sterile 0.15M NaCl, for 5 minutes. Finally, 20ml of medium was added to the alginate beads and cultured in an upright T25 flask at 37°C under 5% CO_2_, for up to 14 days prior to treatment.

### CellTiter-Glo® 3D cell viability assay

For ATP measurement, alginate spheres were seeded in white 96-well plates with one alginate bead per well and treated with each ferroptosis inducer (RSL3, Erastin, FIN56), alongside a 0.2% (v/v) DMSO vehicle control. Treated cells were incubated at 37°C with 5% CO_2_, for 48 hours. After treatments, 100 µl of CellTiter-Glo® reagent from CellTiter-Glo® 3D cell viability assay kit (Promega-UK) was added to each well and mixed for 5 minutes on a plate shaker and incubated at room temperature (RT) for 25 minutes. The luminescent signal was measured using a CLARIOstar plate reader. The average from at least 4 luminescence measurements was calculated and all treated cells were normalized to the controls.

### Immunocytochemistry to detect Nrf2

Untreated MCF-7 and MDA-MB-231 cells were cultured at 2×10^4^ cell per well on 8-well glass chamber slides (Thermo Scientific) for 24 hours prior to immunostaining with1:200 primary Nrf2 antibody (Nrf2 (D1Z9C) XP^®^) rabbit mAb (Cell Signalling Technology, Danvers, USA and 1:500 anti-rabbit IgG (H+L) F(ab’)2 Fragment (Alexa Fluor® 488 conjugate) secondary antibody (Cell Signalling Technology) with a mounted 4,6-diamidino-2-phenyindole (DAPI) (Sigma) counterstain and imaged using a Zeiss LMS 800 confocal microscope.

### Colony formation assays

Radiation exposure of MDA-MB-231 and MCF-7 cells at 1×10^5^ cells/ml at doses of 0, 0.3, 0.6, 1.25, 2.5, 5, 10, 20 Gy was followed by a colony formation assay. Following irradiation, cells were seeded at a density of 1000 cells/ml in 6-well plates and grown in complete medium for 12-15 days. In some experiments, cells were treated with ferroptosis modulators for 24 hours after plating, followed by wash-out for drugs. After 15 days cells were fixed and stained with 0.5% (w/v) crystal violet in dH_2_O for 30 minutes. Colonies containing more than 50 individual cells are counted as an individual colony.

## Results

### The effects of ferroptosis inducers and inhibitors on breast cancer cells

The effects of ferroptosis inducers on cell death of MB-MDA-231 cells and MCF-7 cells was detected using fluorescent microscopy images after staining with Hoechst 33342 and PI (Figure 1A). The ferroptosis inducer RSL3 induced significant cell death at all doses tested from 0.075 µM in MDA-MB-231 cells with greater than 50% cell death at 0.075 µM (Figure 1B). For subsequent combination studies, the optimum dose for combination studies was determined to be 0.0375 µM. MDA-MB-231 cells were highly sensitive to Erastin, showing significant cells death at all doses above 0.15 µM (Figure 1C). Therefore 0.15 µM was considered a suitable dose for combination studies. Propidium iodide-positive cells did not appear to be apoptotic in morphology with absence of pycnotic and condensed nuclei, although this was not formally quantified. The ferroptosis inducer FIN56 induced significant cell death at all doses tested from 0.15 µM, (Figure 1D). Since this dose induced a small, but significant amount of cell death (15%), this was appropriate for use in combination studies with radiotherapy. RSL3 (10 µM) significantly and potently induced cell death in MDA-MB-231 cells (Figure 1E). To establish whether this cell death was likely to be ferroptosis, co-incubation with three ferroptosis inhibitors with differing mechanisms of action was tested, which was completely or partially reversed by ferroptosis inhibitor Deferoxamine, Liproxstatin-1, Ferrostatin-1 (Figure E). Similar results were observed for Erastin and FIN56 (data not shown).

**Figure 1:**
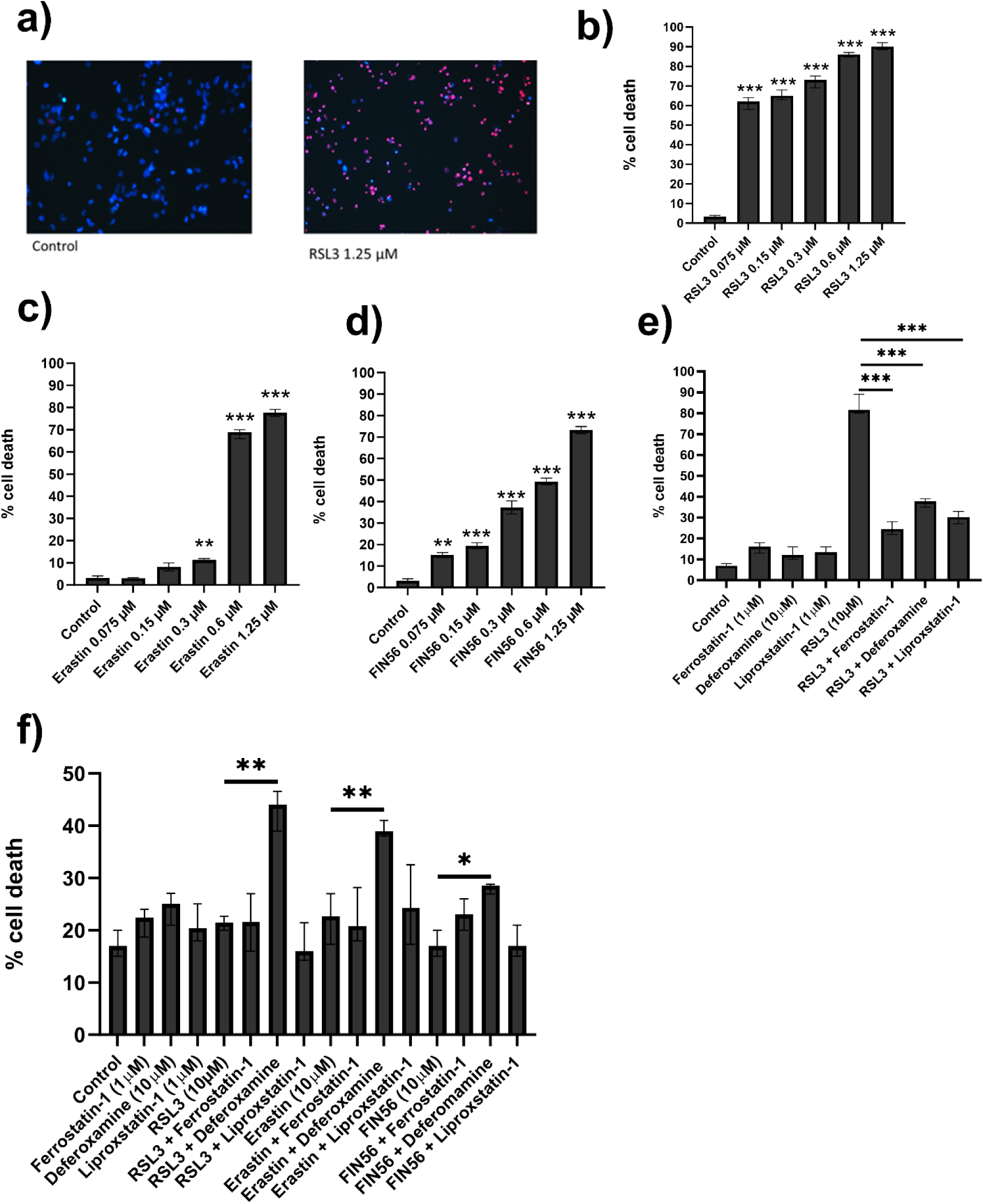
Effect of the ferroptosis inducers RSL3, Erastin and FIN56 and ferroptosis inhibitors on MDA-MB-231 and MCF-7 cells. **(A)** Fluorescence microscopy analysis of cell death in MDA-MB-231 by Hoechst 33342/PI staining after treatment with the ferroptosis inducer RSL3. **(B-D).** Percentage of cell death calculated from 3 independent experiments each analysing triplicate wells for RSL3, Erastin and FIN56 respectively. **(E)** Reversal of ferroptosis after treatment with the ferroptosis inducer RSL3 (10 µM) +/− ferroptosis inhibitors (Deferoxamine 10 µM, Liproxstatin-1 1µM and Ferrostatin-1 1µM) in MDA-MB-231. **(F)** Lack of ferroptosis in response of 10 µM Deferoxamine, Liproxstatin or Ferrostatin, but a significant induction of cells death with deferoxamine in MCF-7 cells. Data is presented as median ± range. The statistical significance was determined by comparison with the control (0.2% (v/v) DMSO), analysed by a Kruskal-Wallis followed by Dunn’s multiple comparisons test (*=P≤0.05, **=P≤0.01, and ***=P≤0.001).

### The effect of ferroptosis inducers and inhibitors on MCF-7 cells

The ferroptosis inducers RSL3, Erastin and FIN56 at 10 µM showed no potent ferroptotic effect on MCF-7 cell death when compared to cells treated with the DMSO vehicle control (Figure 1F). A 10 µM dose of each ferroptosis inducers was the highest dose that can reasonably be added to cells due its solubility in DMSO, so 10 µM was used for all further experiments with MCF-7. The ferroptosis inhibitor Deferoxamine significantly induced cell death when combined with the ferroptosis inducer RSL3 (P≤0.01). This is consistent previous observations specifically in MCF-7 and not MDA-MB-2312 cells in response to Deferoxamine,^22^ however all other ferroptosis inhibitors had little or no effect on MCF7 breast cancer cells.

### The effect of ferroptosis inducers on radiotherapy responses in MDA-MB-231 in 2D cell culture

To assess ferroptosis inducer-induced radiosensitivity, cells were treated with RSL3, Erastin or FIN56 immediately after irradiation with 1.25 Gy and stained with Hoechst 33342 and PI to assess cell death (Figure 2). Although colony formation assay showed potent effects of 1.25 Gy after 10-14 days (see Figure 8), at shorter timepoints used for Hoechst 33342/PI staining, 1.25 Gy had only very modest effects on cell death, so this dose was also used during short-term irradiation studied. Individual treatments did not significantly increase apoptosis or necrosis, but combined treatment significantly increased cell death compared to untreated control cells (P≤0.05), causing an additive effect with Erastin and FIN56 (Figure 2B-C). The data for cell death analysis using ferroptosis inducers-alone was consistent with the results obtained for ATP levels measured using 2D CellTiter-Glo® luminescent cell viability assay (Figure 2A-C), which showed a significant decrease in cell ATP levels after RSL3, Erastin, FIN56 or combination treatment vs. control. However almost all of these effects in combination treatments were due to the ferroptosis inducers. Erastin, RSL3 and FIN56 alone, caused a significant decrease in ATP levels when compared to control cells (P≤0.001). Importantly the ferroptosis inducer and irradiation combined treatment was not significant different from the use of ferroptosis inducer alone. This suggests that the majority of reduction in cell ATP levels in combination treatment was due to ferroptosis inducers, and there was no synergistic enhancement by irradiation (Figure 2).

**Figure 2.**
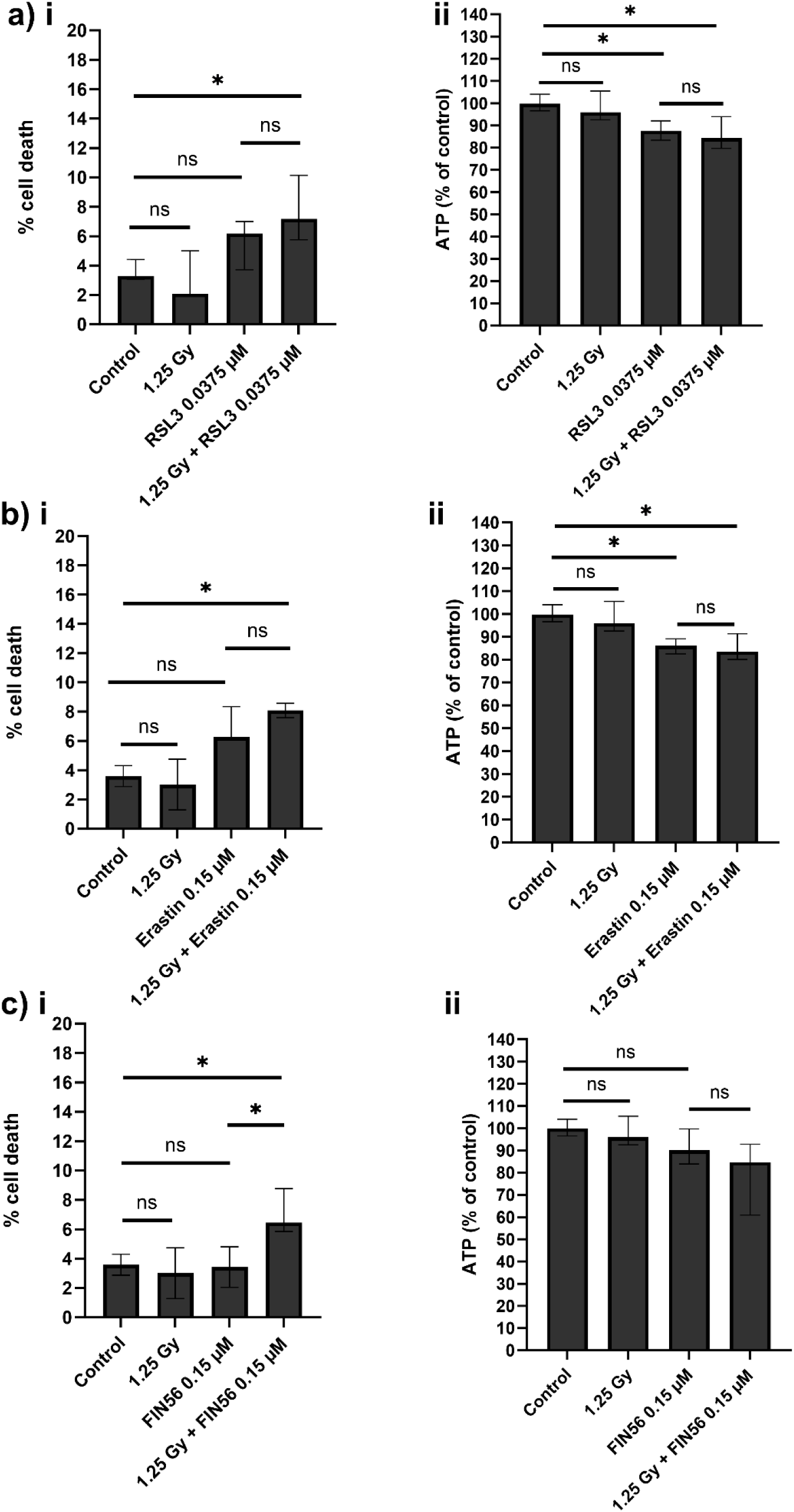
The effect of ferroptosis inducers on radiotherapy responses in MDA-MB-231 in 2D cell culture. MDA-MB-231 cell line were treated with 1.25 Gy of radiation and the ferroptosis inducers **(A)** RSL3 (0.0375 µM, **(B)** Erastin (0.15 µM) or **(C)** FIN56 (0.15 µM) for 72 hours. Assessment of cell death following Hoechst 33342/PI staining was performed and data expressed as percentage of cell death (i), and assessment was made of cell ATP levels using the CellTiter-Glo® luminescent cell viability assay (ii). Data was normalised to the vehicle control which was assigned 100% ATP levels. Data is presented as median ± range. All treatment were repeated in triplicate and repeated in three technical repeats (n=3). The statistical significance was determined by comparison with the control (0.2% (v/v) DMSO), analysed by a Kruskal-Wallis followed by Dunn’s multiple comparisons test (*=P≤0.05, **=P≤0.01, and ***=P≤0.001).

### The effect of ferroptosis inducers on radiotherapy responses in MCF-7 in 2D cell culture

MCF-7 cells were treated with RSL3, Erastin or FIN56 for 72 hours immediately after irradiation and stained with Hoechst 33342 and PI to assess the percentage of cell death and apoptosis and ATP levels (Figure 3). RSL3, Erastin or FIN56 did not synergistically enhance responses to irradiation and combination treatments were not significantly different from RSL3, Erastin or FIN56-alone treatments (Figure 3).

**Figure 3.**
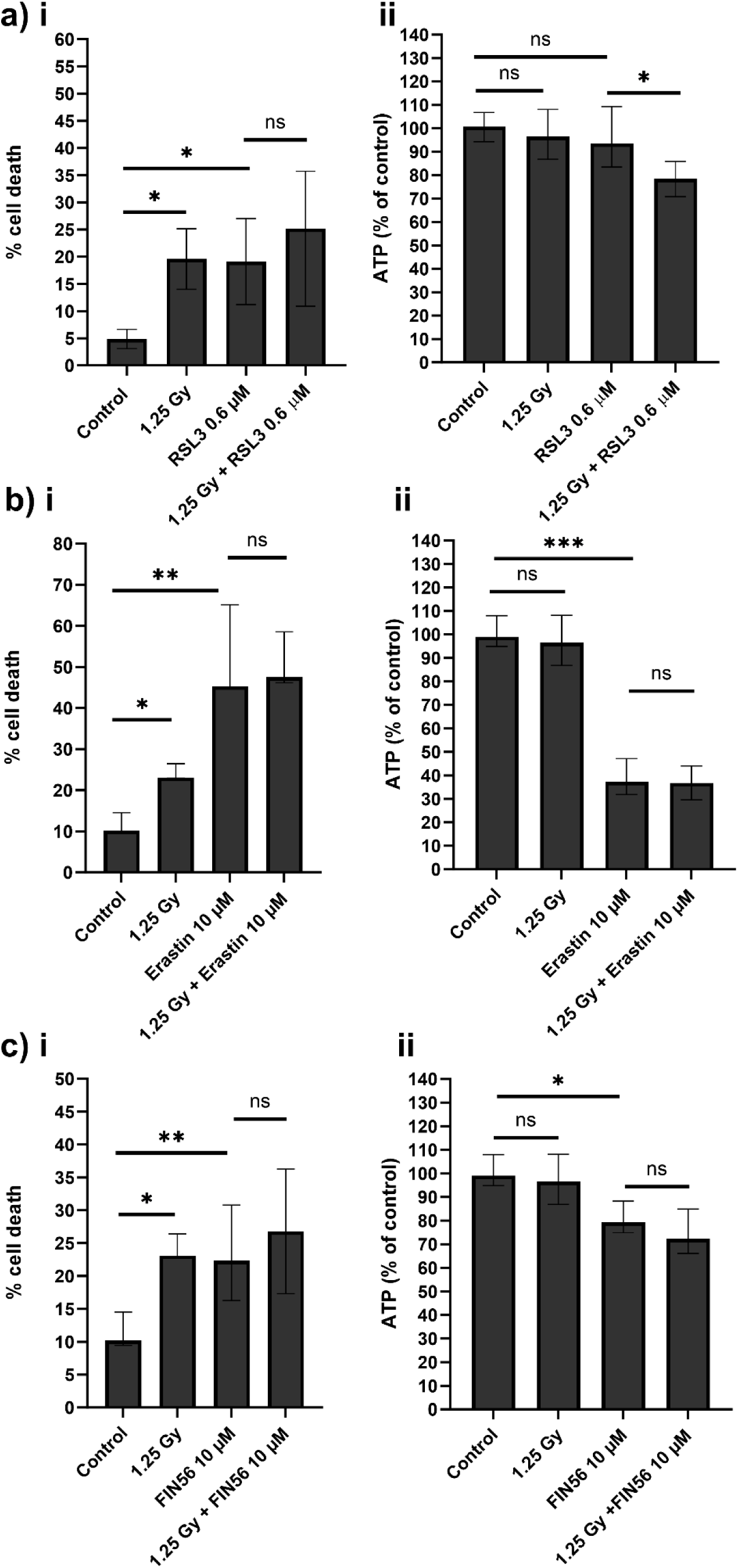
The effect of ferroptosis inducers on radiotherapy responses in MCF-7 in 2D cell culture. MCF-7 cells were treated with 1.25 Gy of radiation and the ferroptosis inducers **(A)** RSL3 (0.0375 µM), **(B)** Erastin (0.15 µM) or **(C)** FIN56 (0.15 µM) for 72 hours. Assessment of cell death following Hoechst 33342/PI staining was performed and data expressed as percentage of cell death (i) and assessment was made of cell ATP levels using the CellTiter-Glo® luminescent cell viability assay. Data was compared to the vehicle control (0.1% v/v DMSO) which was assigned 100% ATP levels. Data is presented as median ± range. All treatment were repeated in triplicate and repeated in three technical repeats (n=3). The statistical significance was determined by comparison with the control (0.2% (v/v) DMSO), analysed by a Kruskal-Wallis followed by Dunn’s multiple comparisons test (*=P≤0.05, **=P≤0.01, and ***=P≤0.001).

### Effect of ferroptosis inducer RSL3 on radiotherapy responses in breast cancer 3D alginate spheroid cells

To assess RSL3-induced radiosensitivity, 3D alginate spheroids were treated with RSL3 for 72 hours immediately after irradiation with 20 Gy and stained with Hoechst 33342 and PI to assess cell death and apoptosis in MDA-MB-231 and MCF-7 cells (Figure 4 and 5 respectively). 20 Gy was chosen, as in parallel studies, 20 Gy induced cell death in some cell lines grown in alginate spheroids after 24 hours, whereas no cell death was observed at doses below 20 Gy (data not shown). Individual treatments (20 Gy in MDA-MB-231, 10 µM RSL3 in MCF-7) did show significant increases in cell death, but effects were additive in combination studies. Here the Hoechst 33342 and PI staining showed different responses of treatments within the 3D spheroids, which clearly show the heterogeneity of responses within populations, in that some spheroids were dead, but others were unaffected (Figure 4 and 5 respectively). Combination treatment showed no enhancement of radiotherapy responses, despite single treatments of 20Gy being used; a dose that normally would prevent formation of all colonies after 14 days in a colony formation assay.

**Figure 4.**
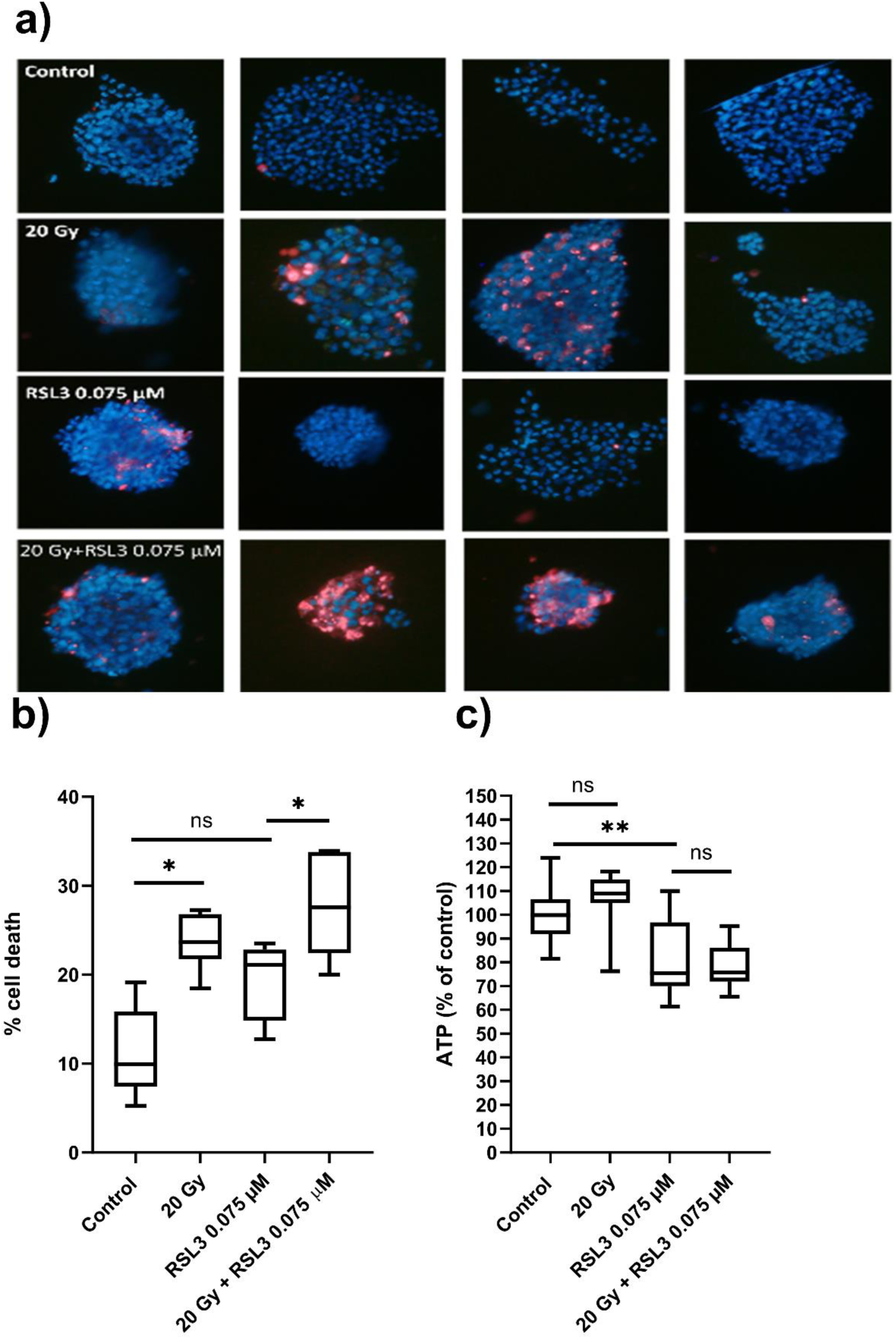
Combination treatment of RSL3 +/− radiotherapy in MDA-MB-231 3D alginate spheroids. **(A)** Hoechst 33342/PI stained MDA-MB-231 spheroids treated with the vehicle control (0.2% (v/v) DMSO), 20 Gy radiation with and without the ferroptosis inducer RSL3 (0.075 µM). **(B)** Cell death count determined following Hoechst 33342/PI staining of MDA-MB-231 spheroids with radiation (20 Gy) and ferroptosis inducer RSL3 (0.075 µM), with a 0.2% (v/v) DMSO vehicle control. Data expressed as median ± range. **(C)** Percentage ATP levels of MDA-MB-231 spheroids level assessed by Cell Titer-Glo® 3D cell viability assay, following treating with radiation (20 Gy) and ferroptosis inducer RSL3 (0.075 µM). ATP levels were expressed as a percentage of the vehicle control (0.2% (v/v) DMSO) which was assigned a 100%. Data expressed as median ± interquartile range. All treatments were repeated in triplicate, in 4 or more technical repeats. The statistical significance was determined by comparison with the control (0.2% (v/v) DMSO), analysed by a Kruskal-Wallis followed by Dunn’s multiple comparisons test (*=P≤0.05, **=P≤0.01, and ***=P≤0.001).

**Figure 5.**
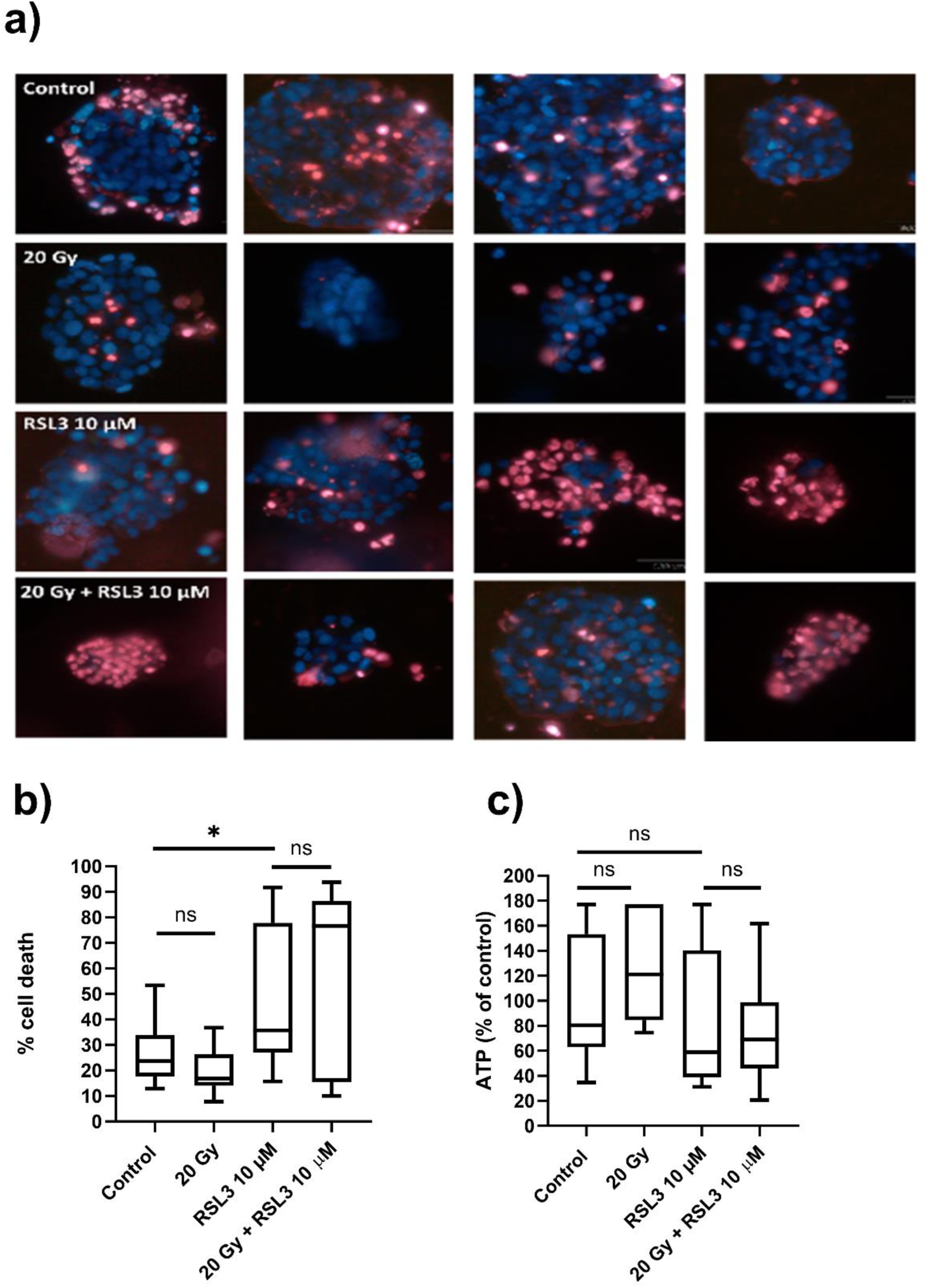
Combination treatment of RSL3 +/− radiotherapy in MCF-7 3D alginate spheroids. **(A)** Cell death was detection by fluorescent microscopy determined by Hoechst 33342/PI staining after treating MCF-7 spheroids with 20 Gy radiation, with control (0.2% (v/v) DMSO) +/− ferroptosis inducer RSL3 (10 µM). Median and interquartile ranges are shown in **(B). (C)** ATP level (% of control) assessed by Cell Titer-Glo® 3D Cell Viability Assay after treating MCF-7 spheroids with radiation (20 Gy) and ferroptosis inducer RSL3 (10 µM), with control (0.2% (v/v) DMSO). Data expressed as median ± interquartile range from n=3 independent experiments each with ≥4 technical repeats. The statistical significance was determined by comparison with the control (0.2% (v/v) DMSO), analysed by a Kruskal-Wallis followed by Dunn’s multiple comparisons test (*=P≤0.05, **=P≤0.01, and ***=P≤0.001).

### The effect of combination treatment of Nrf2 inhibitor ML385 alone and in combination with radiotherapy in breast cancer cells

The basal levels of Nrf2 protein were assessed by immunocytochemistry. In MDA-MB-231, the Nrf2 was very weakly staining in the nucleus (Figure 6A). In MCF-7 cells, Nrf2 was seen more strongly in the nucleus. To assess ML385-inhibitor mediated radiosensitivity, MDA-MB-231 and MCF-7 breast cancer cells were treated with ML385 for 72 hours immediately after irradiation and stained with Hoechst 33342 and PI to assess cell death and apoptosis (Figure 6B-C). ML385 alone significantly increases cell death (P≤0.05), and combined treatments did not further enhance cell death in either cell line. These observations were mirrored by assessment of ATP levels. Therefore, cell death and reductions in cell ATP levels following ML385 treatment was not enhanced by irradiation (Figure 6B-C).

**Figure 6.**
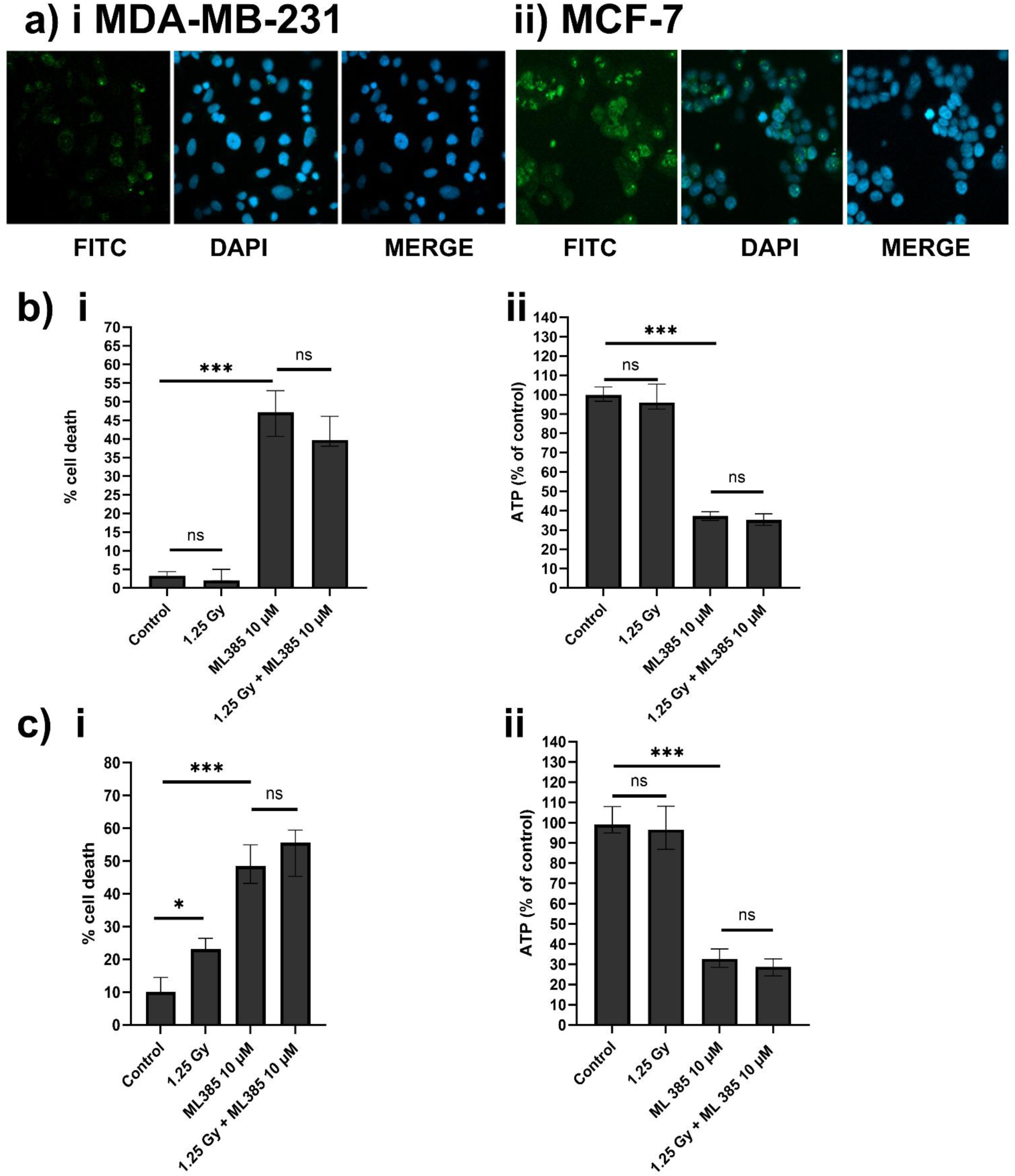
The effect of combination treatment of Nrf2 inhibitor ML385 +/− radiotherapy in MDA-MB-231 and MCF-7 cells. **A)** MDA-MB-231 and MCF-7 cells were stained for Nrf2 using the anti-human Nrf-2 antibody diluted visualized with an AlexaFluor 488-conjugated secondary antibody. Nuclei were counter stained with DAPI. Images were photographed using confocal fluorescence microscopy. Cell death was determined by Hoechst 33342/PI staining under fluorescent microscopy after treating **B)** MDA-MB-231 and **C)** MCF-7 cells with 1.25 Gy radiation, +/− ML385 (10 µM). ATP levels (% of control) were assessed by CellTiter-Glo® luminescent cell viability assay. All treatment were repeated in triplicate and repeated in three technical repeats (n=3). The statistical significance was determined by comparison with the control (0.2% (v/v) DMSO), analysed by a Kruskal-Wallis followed by Dunn’s multiple comparisons test (*=P≤0.05, **=P≤0.01, and ***=P≤0.001).

### Effect of Nrf2 inhibitor on radiotherapy responses in breast cancer 3D alginate spheroid cells

To assess the therapeutic potential of the Nrf2 inhibitor ML385 in spheroids, 3D alginate spheroids were treated with ML385 for 72 hours immediately after irradiation, they were then stained with Hoechst 33342 and PI to assess cell death (Figure 7AB). ML385 treatment alone significantly increased cell death (P≤0.05), but there was no significant increase in cell death when radiation treatment was and combined with ML385. Assessment of cell ATP levels mirrored cell death data, and there was no enhancement of cell death by combination of ML385 with irradiation (Figure 7C).

**Figure 7.**
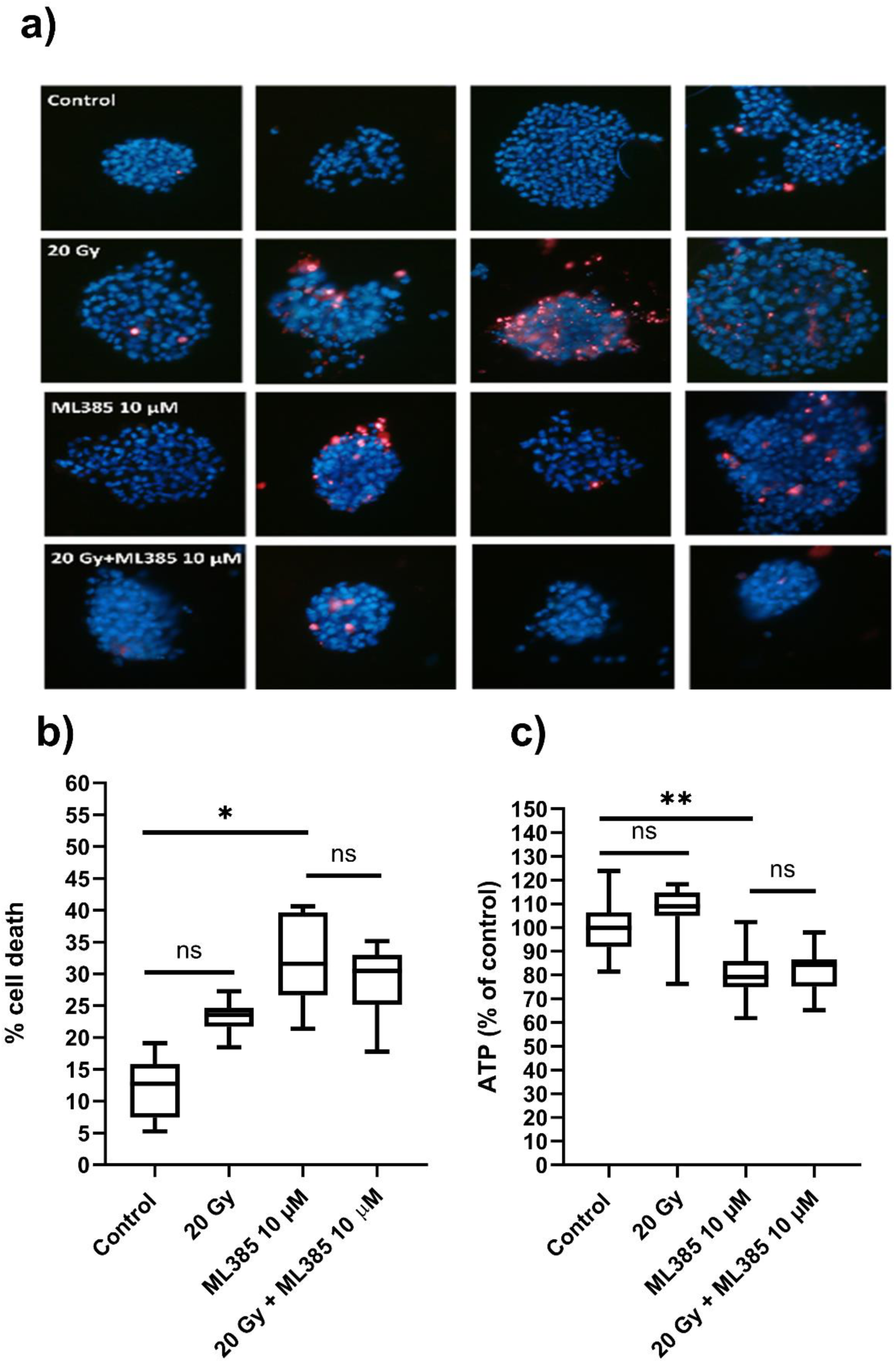
Combination treatment of ML385 +/− radiotherapy in MDA-MB-231 3D alginate spheroids. **(A)** Cell death was determined by Hoechst 33342/PI staining after treating MDA-MB-231 spheroids with 20 Gy radiation +/− the Nrf2 inhibitor ML385 (10 µM), **(B)** Cell death count determined by Hoechst 33342/PI staining after treating MDA-MB-231 spheroids with radiation (20 Gy) and ML385 (10 µM). Data expressed as median ± range. **(C)** ATP level (% of control) assessed by Cell Titer-Glo® 3D cell assay after treating MDA-MB-231 spheroids with radiation (20 Gy) and ML385 (10 µM), with control (0.2% (v/v) DMSO). Data expressed as median ± interquartile range from n=3 independent experiments each with ≥4 technical repeats. The statistical significance was determined by comparison with the control (0.2% (v/v) DMSO), analysed by a Kruskal-Wallis followed by Dunn’s multiple comparisons test (*=P≤0.05, **=P≤0.01, and ***=P≤0.001).

### Effect of ferroptosis inducers and irradiation on colony formation in MDA-MB-231 and MCF-7 cells

Low dose of irradiation between 0 to 2.5 Gy were performed on the colony formation of MDA-MB-231 and MCF-7 cells and the percentage of colony survival was determined (Figure 8A). For both cell lines, 1.25 Gy resulted approximately 60% reduction in colony formation over a 10–14-day period. Since ferroptosis inducers and ML385 showed some cytotoxic effects in standard cell culture and also in 3D cell culture using cell death assays, colony formation assays were performed to assess whether combination treatment affected the number of colonies formed 12-15 days after acute treatment, rather than assessing acute cell death. Consistent with cell death assays, colony formation assays showed cytotoxic effects of 1.25 Gy irradiation, RSL3 and ML385 alone. However, irradiation partially reversed the effects of RSL3, and to a lesser extent Erastin in MDA-MB-231 (Figure 8B). Consistent with cell death studies, MCF-7 showed limited response to RSL3, Erastin or ML385, and combination treatment with 1.25 Gy irradiation was not statistically different to radiation-alone treatment groups (Figure 8B).

**Figure 8.**
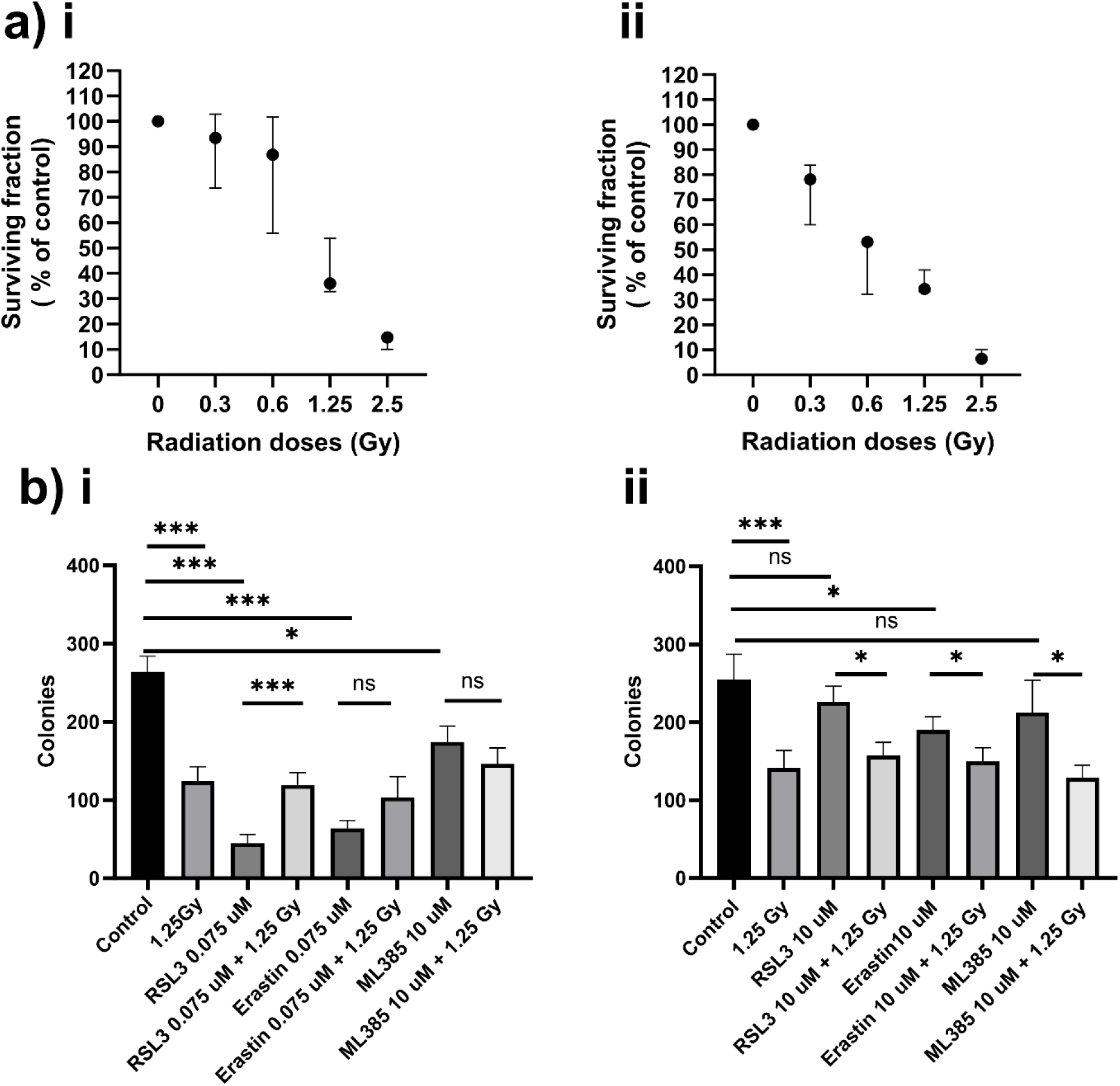
Colony formation assay following irradiation and treatment with ferroptosis inducers. **A)** Colony formation of i) MDA-MB-231 and ii) MCF-7 breast cancer cells grown for 12-15 days following irradiation at 0 to 2.5 Gy cells culture and stained with crystal violet and the number of colonies formed were counted and compared to untreated control cells which were assigned a 100% survival rate. All treatment were repeated in triplicate and repeated in three technical repeats (n=3). **B)** Cells were irradiated and/or treated with ferroptosis modulators followed plating at clonal density. Data expressed as median ± range from n=3 independent experiments each with 3 technical repeats. The statistical significance was determined by comparison with the control (0.2% (v/v) DMSO), analysed by a Kruskal-Wallis followed by Dunn’s multiple comparisons test (*=P≤0.05, **=P≤0.01, and ***=P≤0.001).

## Discussion

The aim of this study was to assess whether radiotherapy responses could be enhanced in 2D and 3D cell cultures of breast cancer cells with modulators of ferroptosis signaling. To assess this, MDA-MB-231 and MCF-7 cells were treated with ferroptosis inducers and Nrf2 inhibitors after radiotherapy and assessed by direct assessment of cell death, and by measurements of ATP as a marker of cell viability. In summary, in these studies, MCF-7 and MDA-MB-231 breast cancer cells do not show enhanced radiosensitivity when treated with either ferroptosis inducers, or a Nrf2 inhibitor.

Colony formation assays were performed on MDA-MB-231 and MCF-7 cells to initially optimize the irradiation dose (Figure 8). 1.25 Gy reduced colony formation by approx. 60%, and this dose was also used for cell death experiments. The rationale was that colony formation showed reduced numbers of cells able to proliferate over 2 weeks, in-part in response to DNA repair being successfully resolved, so 1.25 Gy must be affecting cells in short-term studies also.

Both MCF-7 and MDA-MB-231 were irradiated and then immediately treated with doses of RSL3, Erastin or FIN56 at doses that were known to induce cell death in a minority of cells but not induce high levels of cell death (Figure 2). In all cases, there was no consistent synergistic response in combination treatment. In only FIN56 treatment of MDA-MB-231 showed evidence of a weak synergistic effect when combined with irradiation (Figure 2C). In all cases, ATP levels generally followed cell death observations with no synergistic responses seen.

There is evidence to suggest that ferroptosis inducers such as Erastin may enhance the sensitivity of cancer cells to radiotherapy.^14^ Erastin inhibits system Xc-reducing cysteine import and hence decreased glutathione, meaning there is less glutathione to respond to irradiation-induced ROS. In addition, some studies have suggested that ferroptosis may play a role in the radiosensitivity of cancer stem cells ^13 23^ which are thought to be responsible for cancer recurrence and resistance to radiotherapy. On the other hand, cancer cells after radiation may evade cell death via modulation of ferroptosis by several mechanisms. In the presence of a Keap-1 mutation, Nrf2 is high, leading to an enhanced antioxidant response. Irradiation-mediated ROS may have little effect in this situation. Similarly, Acyl coenzyme A synthetase long chain family member 4 (ACLS4) is low in some breast cancers, this is primarily responsible for catalysing the conversion of free PUFAs such arachidonic acids (AAs) and adrenic acids (AdAs) to their acyl coenzyme A (CoA) derivatives, such as AA/AdA-CoA. These PUFA-CoAs are then converted into lysophospholipids (LysoPLs), which are then further incorporated into phospholipids (such AA-PE and AdA-PE) by lysophosphatidylcholine acyltransferase 3 (LPCAT3) and other enzymes. Hence, PUFA-PL synthesis is suppressed and ferroptosis resistance is markedly increased when ACSL4 or LPCAT3 are suppressed.^312^ This was supported by other studies of radioresistant sublines of MCF-7 cells, which were showed a loss of ACSL4. ^24^ Furthermore, ASCL4 is essential for mediating radiation-induced damage in normal tissues, and specific ASCL4 inhibitors are being trialled as inhibitors of ferroptosis-induced radiation damage.^25^

The major ferroptosis defence included the solute carrier family 7 member 11-glutathione-GPX4 (SLC7A11-GSH-GPX4).^1^ A crucial part of the cystine/glutamate antiporter system Xc^−^, SLC7A11, facilitates the antiporter action of system Xc^−^ by bringing in extracellular cystine and releasing intracellular glutamate.^26^ Irradiation is known to increase SLC7A11 expression, leading to increased cystine uptake and increased GSH production, resulting in a reduced response to radiation,^26^ and therefore SLC11A7 overexpression leads to radio-resistance,^27,28^ whereas decreased SLC11A7 leads to radiosensitivity.^29^ After extracellular cystine is taken up by SLC7A11, it is immediately reduced to cysteine in the cytosol through a reduction mechanism that consumes NADPH. Next, as a key cofactor for GPX4 to detoxify lipid peroxides, cysteine acts as the rate-limiting precursor for the manufacture of GSH.^30^Many cancer cells undergo significant ferroptosis when SLC7A11 transporter activity is blocked or when cystine is not present in culture media. ^30^ Importantly, some tumour suppressors, including p53, BAP1, ADP-ribosylation factor (ARF), and Kelch-like ECH-associated protein 1 (KEAP1), induce ferroptosis by inhibiting the production or function of SLC7A11. ^31 31^. Similarly, by binding to the SLC7A11 promoter, activating transcription factor 3 (ATF3) suppresses SLC7A11 expression and increases the vulnerability of cancer cells to ferroptosis.^32^ Nrf2 and activating transcription factor 4 (ATF4) are examples of stress-responsive transcription factors that can promote SLC7A11 under a variety of stress circumstances, including oxidative stress and amino acid deficiency, protecting cells from ferroptosis. ^33^. Therefore, SLC7A11 is an endogenous protective protein against radiation induced damage in normal cells, which can be exploited immediately after radiotherapy by tumour cells and can be over-expressed in tumour cells to generate an intrinsically radioresistant phenotype, potentially contributing to an absence of radio-sensitization my ferroptosis and Nrf2 modulators in this study.

The Nrf2 inhibitor, ML385 was combined 1.25 Gy for 2D and 20 Gy for 3D cell showed no enhancement of radiation responses in both MDA-MB-231 and MCF-7. Since ML385 is a direct inhibitor of Nrf2, it was hypothesised that ML385 would attenuate the Nrf2-mediated antioxidant response which becomes activated after irradiation and increases cell death. Despite Nrf2 expression being much higher in MCF-7 cells than MDA-MB-231 (Figure 6), both cell lines responded similarly to ML385 alone suggesting that in standard growth conditions, both are dependent on Nrf2-mediated antioxidant response to survive. In both cases, combination of ML385 ± irradiation resulted in essentially no difference in cell death, either by ATP measurements or Hoechst 33342/PI staining. The reason for this is unclear although cells were treated with ML385 within 2 hours of irradiation which may be a factor. Further experiments with pre-treatment are required. However, ML385 sensitises other cancer cells to irradiation in other tumour models, such as lung cancer. ^34^ Furthermore, the formation of ROS and DNA damage are two factors that affect radiotherapy effectiveness. Excessive ROS production can cause cell death or activate defence mechanisms like the Keap1/Nrf2 pathway, which controls intracellular cysteine availability by upregulating SLC7A11, and subsequently glutathione synthesis, enhancing antioxidative defence. ^34^

Cells grown in 3D alginate spheres showed limited radiosensitivity at 20 Gy, whereas at this dose, almost all cells were killed in 2D colony formation assays (Figure 4-5 and Figure 8). It is known that cells in 3D cell culture experience increased ROS, and induce Nrf2 in the hypoxic core, and that Nrf2 expression is a pre-requisite for sphere formation.^17^ Ionizing radiation damages DNA directly or indirectly, producing ROS and causing additional oxidative stress-related damage to biomolecules. Tumour cells evolve defence systems to prevent cell death due to repeated ROS exposure, and gain of Nrf2, or in the case of some breast cancers loss of Keap-1, allows clonal evolution of surviving Nrf2-high cells.^17^ These Nrf2-dependent mechanisms enhance DNA repair and boost anti-oxidation defence, neutralising ROS, reducing oxidative stress, and limiting ROS-induced damage. ^35^ Furthermore hypoxia-inducible factors (HIFs) in the hypoxic tumour microenvironment are a significant radio-resistance mechanism. ^36^ This occurs in 3D cell culture as a mimic of *in vivo* responses.

Cells grown in 3D cell culture and treated with the ferroptosis inducer RSL3 (Figure 4-5) and to some extent ML385 (Figure 7) showed that some colonies respond to treatment and die, while other colonies survive showing heterogeneity within these populations of tumour cells. Interestingly, a parallel phenomenon has very recently been described in TNBC whereby cancers show marked heterogeneity with respect to ferroptosis markers. ^37^ Similarly, ferroptotic heterogeneity has also been reported in melanoma^38^ which mirrors observations for heterogeneity of apoptosis inducers in breast cancer cells and other tumour cell types. ^39^ Further research is necessary to determine the possibility and precise processes by which ferroptosis activation and Nrf2 inhibition can decrease the radio-resistance of hypoxic cancer cells in various cancer situations, and to identify metabolic signatures of ferroptosis-sensitive vs. resistant populations.

In summary, MDA-MB-231 and MCF-7 cells were sensitive to irradiation in 2D cell culture but resistant to irradiation in 3D cell culture. After induction of cell death with ferroptosis inducers, there was no short-term robust enhancement to radiation effect to the breast cancer cells either in 2D or 3D cell culture. The Nrf2 inhibitor ML385 showed no further effect on irradiation. These studies suggest targeting ferroptosis does not induce short-term enhancement of ferroptotic cell death, or long-term colony formation after acute combination treatment (Figure 8). Further studies using pre-treatment of ferroptosis prior to radiotherapy are required, as well as studies using repeated fractionation of doses over a prolonged period of time to mimic the clinical situation, which is particularly relevant to treatment of 3D spheroids.

## Acknowledgements

This study was funded by a PhD scholarship from the Kuwait Embassy Cultural Office.

## Author contributions

AA completed the majority of the laboratory work and data analysis and edited the manuscript, NAC conceived the study, drafted the manuscript and completed colony formation assays, NJM conceived some of the experiments and supervised the work, CS edited the manuscript and completed confocal microscopy analysis.

## Data availability

The datasets used and/or analyzed during the current study are available from the corresponding author on reasonable request.

## Other information

The authors have no competing interests to declare

## Notes

### Competing Interest Statement

The authors have declared no competing interest.

### Summary of Updates

The revised document has additional data added, specifically colony formation assays in Figure 8. The manuscript has been condensed (combined figures, removed unnecessary images) to allow direct journal submission but all previous key data remains. Some statistical analyses have been improved, and some graphs redrawn for clarity.

